# *Hugin*^+^ neurons link the sleep homeostat to circadian clock neurons

**DOI:** 10.1101/2020.04.29.068627

**Authors:** Jessica E. Schwarz, Anna N. King, Cynthia T. Hsu, Annika F. Barber, Amita Sehgal

## Abstract

Sleep is controlled by homeostatic mechanisms, which drive sleep after wakefulness, and a circadian clock, which confers the 24-hour rhythm of sleep. These processes interact with each other to control the timing of sleep in a daily cycle as well as following sleep deprivation. However, the mechanisms by which they interact are poorly understood. We show here that *hugin*+ neurons, previously identified as neurons that function downstream of the clock to regulate rhythms of locomotor activity, are also targets of the sleep homeostat. Sleep deprivation decreases activity of *hugin*+ neurons, likely to suppress circadian-driven activity during recovery sleep, and manipulations of *hugin*+ neurons affect sleep increases generated by activation of the homeostatic sleep locus, the dorsal fanshaped body (dFB). Also, mutations in peptides produced by the *hugin*+ locus increase recovery sleep following deprivation. Trans-synaptic mapping reveals that *hugin*+ neurons feed-back onto central clock neurons, which also show decreased activity upon sleep loss, in a Hugin-peptide dependent fashion. We propose that *hugin*+ neurons integrate circadian and sleep signals to modulate circadian circuitry and regulate the timing of sleep.

## Introduction

Sleep is regulated by two processes, circadian and homeostatic ^1^. The endogenous circadian clock, together with its downstream pathways, is synchronized to external day-night cycles and determines the timing of sleep to produce 24-hour rhythms in sleep and wake. The homeostatic process tracks sleep:wake history and generates sleep drive based on the extent of wakefulness. Overtly, sleep homeostasis can be seen as an increase in sleep duration and depth after prolonged wakefulness. Generally, circadian and homeostatic processes are studied as separate mechanisms that regulate sleep, but they clearly intersect and are coordinated in a daily cycle to promote the onset and maintenance of sleep at night. Following sleep deprivation, the homeostatic system can drive sleep at the *wrong* time of day, but even under these conditions, interactions between the two systems determine the timing and duration of sleep. However, the neuronal mechanisms by which circadian and homeostatic pathways signal to each other are unknown.

The functions and regulation of sleep are extensively studied in model organisms, such as *Drosophila melanogaster*^2^. In the *Drosophila* brain, the circadian clock is expressed in ~150 clock neurons that are organized into neuroanatomical groups, of which the ventral and dorsal lateral neurons are the most important for driving rhythms of locomotor activity ^3,4^. The small ventrolateral neurons (s-LNvs) link to other brain regions is through different circuits. One of those circuits connects the sLNvs to the site of the motor ganglion, the thoracic nerve cord: s-LNvs → DN1s → *Dh44*+ neurons → *hugin*+ neurons→ ventral nerve cord ^5^. *Dh44*-expressing neurons in the pars intercerebralis regulate locomotor activity rhythms in part through the signaling of DH44 neuropeptide to *hugin*-expressing neurons in the subesophageal zone (SEZ) ^6,7^. *Dh44*+ and *hugin*+ circadian output neurons do not contain canonical molecular clocks, but display cycling in neuronal activity or peptide release, likely under control of upstream circadian signals ^7–9^. Links between these neurons and loci regulating sleep homeostasis have not been identified yet.

Regulation of sleep homeostasis in flies involves the central complex and mushroom body ^10–13^. Recent studies have focused on a group of sleep-promoting neurons that project to the <u>d</u>orsal <u>f</u>an-shaped <u>b</u>ody (dFB) neuropil in the central complex. Activation of dFB neurons labeled by the *23E10-GAL4* driver promotes sleep ^14,15^, and these neurons are required for normal sleep rebound after deprivation ^16^. *23E10*+ neurons receive input signals from R2 ellipsoid body neurons, which track sleep need ^17^.

As *hugin*+ neurons are significantly downstream of central clock neurons and close to behavioral outputs, we asked if they also have a role in sleep. We find that *hugin*+ neurons are dispensable for determining daily sleep amount, but they show decreases in activity following sleep deprivation. They also receive projections from the dFB and counter sleep-promoting effects of *23E10*+ neurons, such that ablation of *hugin*+ neurons enhances sleep driven by *23E10+* cells. Further supporting a role in sleep, activation of *hugin*+ neurons and mutations in Hugin peptides affect recovery sleep after deprivation. *hugin*+ neurons target PDF+ s-LNv clock neurons, which also show decreases in intracellular Ca^2+^ levels following sleep deprivation. Thus *hugin*+ neurons serve as a nexus between the sleep homeostat and the circadian clock.

## Results

### Silencing *hugin*+ neurons does not affect sleep

As noted above, because *hugin*+ neurons mediate circadian output and are just upstream of the motor ganglion^7^, we asked if they also regulate sleep. To test this, we expressed temperature-sensitive *TrpA1* channel ^18^ in *hugin*+ neurons and activated them with high temperature while measuring sleep behavior. In other experiments, we expressed temperature-sensitive *shibire^ts^*, a dominant-negative dynamin gene ^19^, to inhibit synaptic transmission from *hugin*+ neurons at high temperature. Unlike *23E10*+ neurons that increase sleep when activated with TrpA1 ^14,15,20^, neither activation of *hugin*+ neurons with *TrpA1* (Figure S1A) nor inhibition with *shibire^ts^* (Figure S1B) produced any changes in sleep. Despite unchanged sleep levels, *hugin*>*shibire^ts^* flies were less active than control flies, as measured by number of beam crossings per day (Figure S1C), which confirms our previous findings that *hugin*+ neurons regulate locomotor activity ^7^.

Since mechanisms that participate in baseline and sleep recovery may be different, we asked whether *hugin*+ neurons play a role in regulating sleep homeostasis. We used the same thermogenetic approaches to activate or inhibit the *hugin*+ neurons while simultaneously sleep depriving the flies at night using a mechanical method. After sleep deprivation, recovery sleep was monitored in the flies during the daytime. We found no significant difference in recovery sleep between the experimental and control genotypes when *hugin*+ neurons were activated or inhibited (Figure S1D). Given that mechanical sleep deprivation can recruit multiple pathways to elicit rebound ^21^, it is possible that disrupting the activity of *hugin*+ neurons alone does not affect sleep amount or homeostasis.

### Sleep deprivation decreases Ca^2+^ levels in *hugin*+ neurons

Although *hugin*+ neurons did not affect sleep, we considered the possibility that they were affected by sleep loss. Sleep is correlated with changes in neuronal activity in sleep-regulatory circuits, including the MB, dFB, and R2 ellipsoid body ^12,17,22,23^. For example, sleep-promoting dFB neurons tend to be more electrically active after sleep deprivation, when sleep pressure is high, than in rested flies ^24^. To determine if sleep loss alters neuronal activity in *hugin*+ neurons, we measured intracellular Ca+ using CaLexA (Calcium-dependent nuclear import of LexA) ^25^, which drives expression of GFP in response to sustained increases in Ca^2+^ levels. We used *hugin-GAL4* to express CaLexA-GFP transgenes and *UAS-CD8:RFP* for normalizing the GFP signal. *hugin*>*CaLexA-GFP;RFP* flies were deprived of sleep for nine hours at the end of the night (ZT 15-24) and subsequently collected for GFP measurements (Figure 1A). A control group, flies that were not sleep deprived, was assayed at the same time of day as the deprived group. In the sleep-deprived flies, the CaLexA-dependent GFP signal in *hugin*+ neurons was lower than in control flies (Figure 1B-C). To rule out a general effect of sleep deprivation on Ca^2+^, we also tested if sleep deprivation affects Ca^2+^ levels in *Dh44*+ neurons, another group of circadian output neurons ^6^. However, the CaLexA-GFP signal in *Dh44*+ neurons was not significantly different between the sleep-deprived and control flies (Figure 1D). Thus, sleep deprivation specifically affects Ca^2+^ levels of *hugin*+ neurons, suggesting that the homeostat inhibits *hugin*+ neurons.

**Figure 1:**
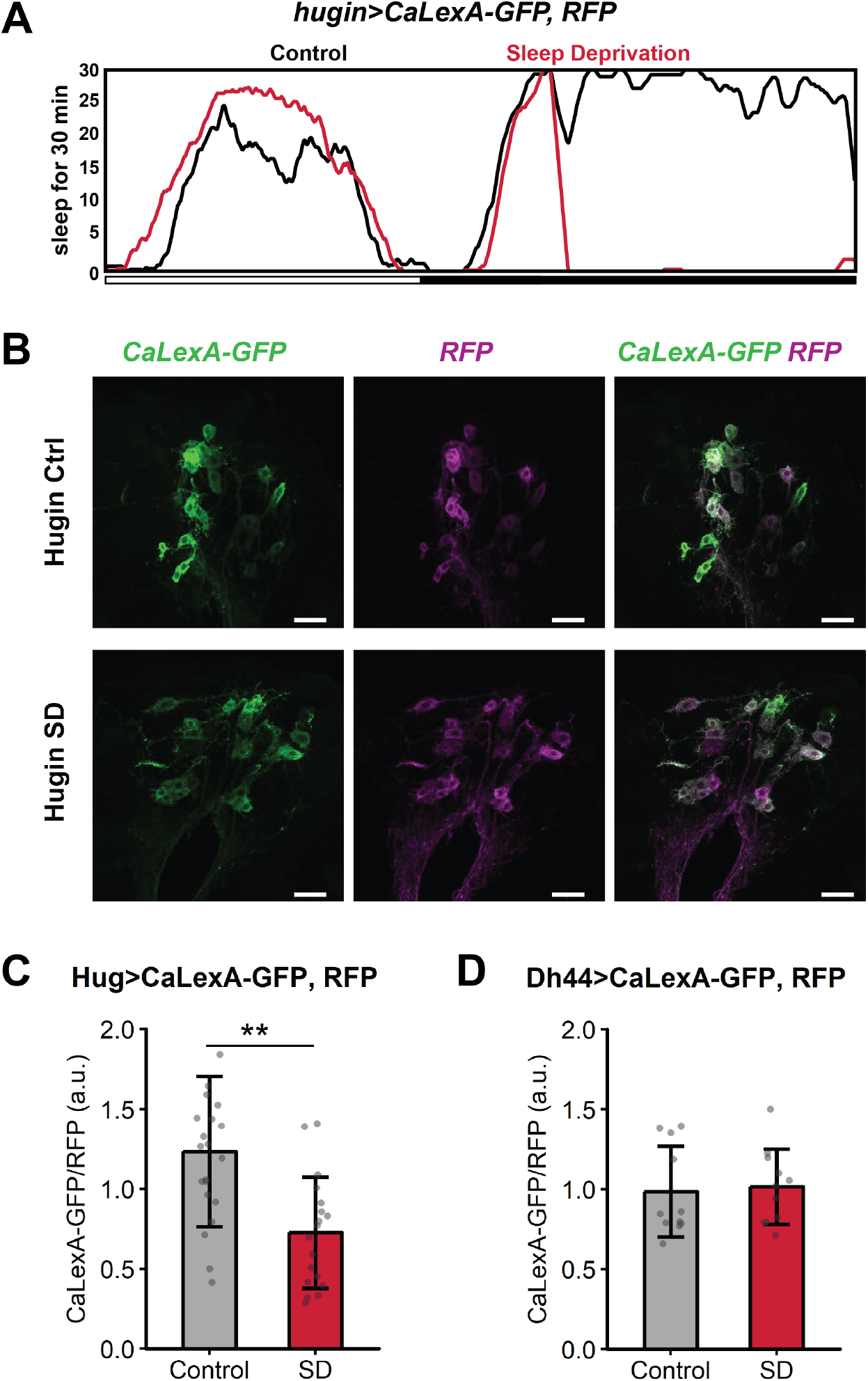
Ca^2+^ levels of *hugin*+ neurons are suppressed with sleep deprivation. **(A)** Sleep profiles of *hugin*>*CaLexA-GFP; RFP* flies subjected to no sleep deprivation (Control, black, n = 8 flies) or 9-hr sleep deprivation (SD, red, *n* = 8 flies). Sleep graphed as minutes per 30-minute bin over 21 hours. **(B)** Representative images show GFP reporting Ca^2+^ levels via the CaLexA system and RFP normalizer signals in *hugin*>*CaLexA-GFP; RFP* flies from Control or SD groups. Max intensity projection images show *hugin*+ neurons in subesophageal zone. Scale bar, 25 μm. **(C)** Levels of GFP signal normalized to RFP signal in *hugin*+ cell bodies from Control (*n* = 21 flies) and SD (*n* = 18 flies) groups. **P = 0.000449, Welch’s *t*-test. **(D)** Levels of GFP signal normalized to RFP signal in *Dh44*+ cell bodies from Control (*n* = 11 flies) and SD (*n* = 11 flies) groups. n.s., P = 0.782 by Welch’s *t*-test.

### *hugin*+ neurons receive projections from the sleep-promoting dFB

Given that *hugin*+ neurons are affected by sleep loss, we sought to determine if they are connected to circuitry of the sleep homeostat. Based upon our previous findings that *hugin*+ neurons project back to the dorsal part of the fly brain ^7^, including the superior medial protocerebrum (SMP) in the vicinity of the dFB, we asked if *hugin*+ neurons contact the dFB. Using the sleep-promoting *23E10-GAL4* driver ^16,24,26,27^ to label dFB neuron dendrites and simultaneously using the LexA system to mark pre-synaptic sites of *hugin*+ neurons, we found that both sets of projections localize to the SMP (Figure S2). Published images also suggest presynaptic sites of dFB neurons are in the SMP, even though presynaptic sites are primarily localized in a single dorsal layer of the fan-shaped body ^20,27–29^. Using *23E10-LexA* to express *Rab3::GFP* we confirmed the presence of presynaptic projections of *23E10*+ neurons in the dFB and SMP, albeit with a weaker signal in the SMP (Figure 2A). Additionally, we expressed *brp-short^GFP^*, a nonfunctional 754-residue portion of bruchpilot (BRP) that localizes to presynaptic active zones ^30,31^, in *23E10*+ neurons. This pre-synaptic marker labeled projections in both the dFB and SMP (Figure 2B).

**Figure 2:**
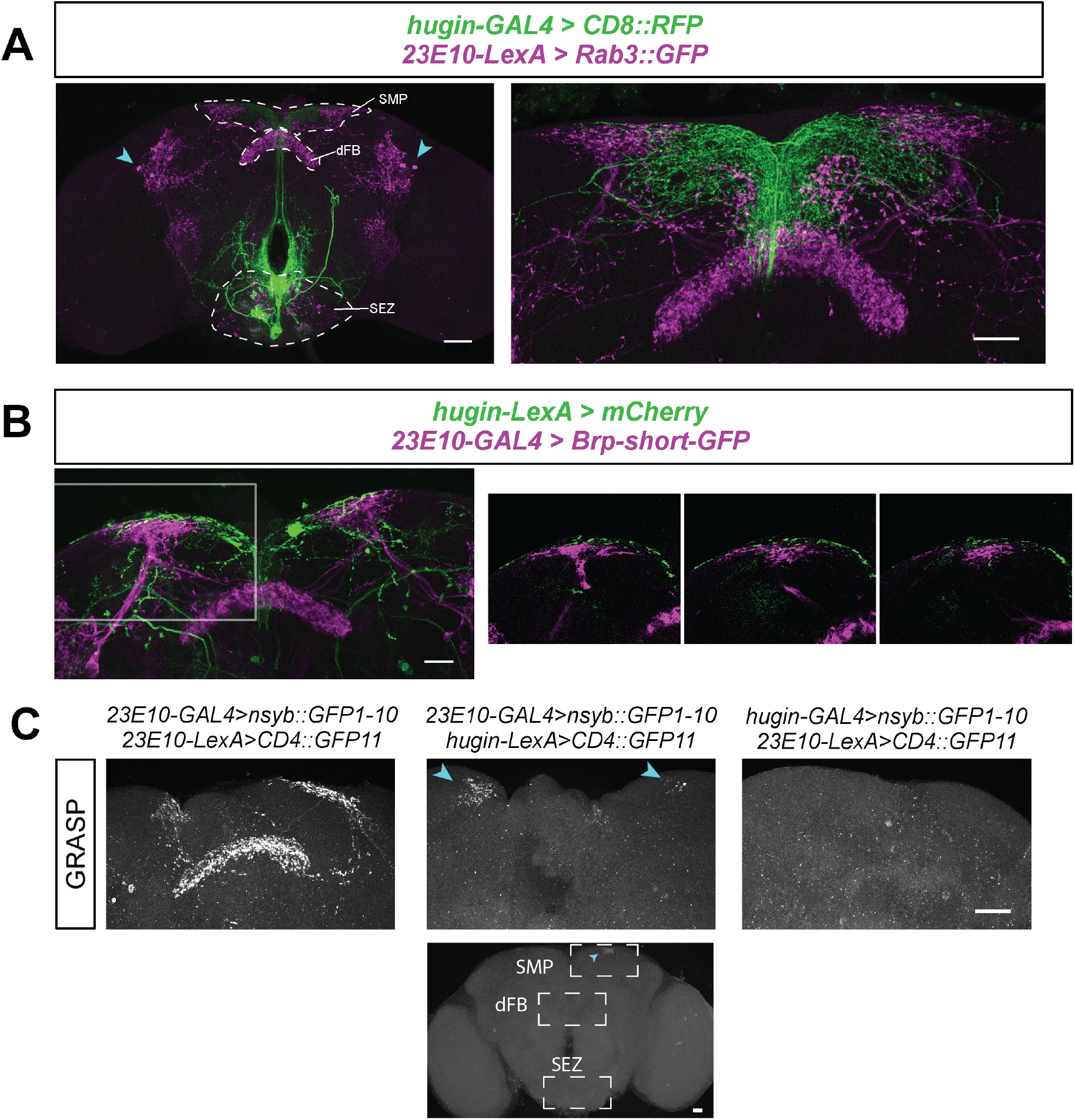
Sleep-promoting dFB (<u>d</u>orsal <u>f</u>an-shaped <u>b</u>ody) neurons contact *hugin*+ circadian output neurons. **(A)** Co-labeling of *hugin*+ neurons with membrane marker (green) and *23E10*+ dFB neurons with RAB3::GFP, a presynaptic marker (magenta). The left image shows co-labeling of neurons in the whole fly brain; arrowheads indicate *23E10*+ cell bodies. Superior medial protocerebrum (SMP), dorsal fan-shaped body (dFB), and subesophageal zone (SEZ) regions are labeled. The right image shows the dorsal protocerebrum, where *hugin*+ projections intermingle with *23E10*+ projections in the SMP. **(B)** Co-labeling of *hugin*+ neurons with membrane marker (green) and *23E10*+ dFB neurons with BRP-short^GFP^, a presynaptic marker (magenta). The left image shows co-labeling of neurons in the dorsal brain. The right series of images show single confocal sections of the region indicated by white box, where *hugin*+ projections intermingle with *23E10*+ projections in the SMP. **(C)** Synaptic nSyb::spGFP1-10 is expressed in presynaptic neurons and complementary spGFP11 expressed in putative postsynaptic neurons. GFP reconstitution occurs only if synaptic connectivity exists. C, Left: When both nSyb::spGFP1-10 and spGFP11 are expressed in *23E10*+ dFB neurons, GFP reconstitution occurs in the dFB and SMP. C, Middle, Top: Cyan arrowheads point to the GFP reconstitution in the SMP when nSyb::spGFP1-10 is expressed in *23E10*+ dFB neurons and spGFP11 is expressed in *hugin*+ neurons. C, Middle, bottom: GFP is only reconstituted in the SMP (not dFB or SEZ) when nSyb::spGFP1-10 is expressed in *23E10*+ dFB neurons and spGFP11 is expressed in *hugin*+ neurons. C, Right: No GFP reconstitution when nSyb::spGFP1-10 is expressed in *hugin*+ neurons and spGFP11 is expressed in *23E10*+ dFB neurons. Scale bars, A(left): 50 μm; A(right), B, C: 25 μm.

To look for a possible synaptic connection between *23E10*+ and *hugin*+ neurons, we used a trans-synaptic GFP fluorescence reconstitution assay (nSyb-GRASP). This system uses the expression of a split GFP, one part tethered to neuronal Synaptobrevin (nSyb::spGFP1-10) in the putative presynaptic cells and the complement tethered to the membrane (CD4::spGFP11) of the putative postsynaptic neurons ^32^. As opposed to the original GRASP that did not label synapses specifically ^33^, trafficking of nSyb to the presynaptic vesicle membrane ensures that nSyb-GRASP identifies membrane contacts specifically at synapses and also indicates directionality of the contact. We first tested the nSyb-GRASP tool by coexpressing both presynaptic nSyb::spGFP1-10 and complementary CD4::spGFP11 in *23E10*+ dFB neurons. In these flies, GFP reconstituted in both the dFB and SMP (Figure 2C left), confirming that *23E10*+ dFB neurons have presynaptic sites in both these regions. As an additional control, we made sure no GFP fluorescence was observed in brains expressing either half of the GRASP components and imaged under the same conditions (data not shown). In flies with the presynaptic nSyb::spGFP1-10 expressed in *23E10*+ dFB neurons and complementary CD4::spGFP11 expressed in *hugin*+ neurons, fluorescent GFP reconstituted in the SMP, but not in the dFB, or in the SEZ (Figure 2C middle, top and bottom). We also performed the reciprocal experiment, with nSyb::spGFP1-10 expressed in the *hugin*+ neurons and complementary CD4::spGFP11 expressed in *23E10*+ dFB neurons, but did not observe any GFP fluorescence in the brain (Figure 2C right), suggesting that *hugin*+ neurons do not signal to dFB cells. As GRASP signals can be difficult to detect, we cannot entirely exclude the possibility of a reciprocal connection, but these data clearly demonstrate that *23E10*+ dFB neurons are presynaptic to *hugin*+ neurons in the SMP.

### *hugin*+ neurons modulate output of *23E10*+ sleep-promoting dFB neurons

We next tested whether *hugin*+ neurons affect the sleep-promoting output of *23E10*+ neurons, by activating *23E10*+ dFB neurons in flies where *hugin*+ neurons are ablated (Figure 3A-B). We confirmed that expression of Reaper in *hugin*+ neurons ablated these neurons by the loss of co-expressing GFP signal (Figure S3A) and by monitoring behavioral phenotypes. Similar to our previous work where we inhibited *hugin*+ neurons^7^, we found that ablation of *hugin*+ neurons significantly decreased morning activity and also resulted in a trend towards lower activity in the evening (ZT6-12) (Figure S3B). Thermogenetic activation of the *23E10*+ neurons using the LexA/LexAop system to drive two copies of TrpA1 (*23E10-LexA*>*LexAop-TrpA1(2x);* +>*UAS-reaper*) led to sleep increase during the day as reported previously ^14,15,20^ (Figure 3A-B). Activation of *23E10*+ neurons in flies lacking *hugin*+ neurons (*23E10-LexA*>*LexAop-TrpA1(2x); hugin-GAL4*>*UAS-reaper*) enhanced the typical daytime increase in sleep (Figure 3B, second graph). This result indicates that *hugin*+ neurons counter the output of *23E10*+ neurons.

**Figure 3:**
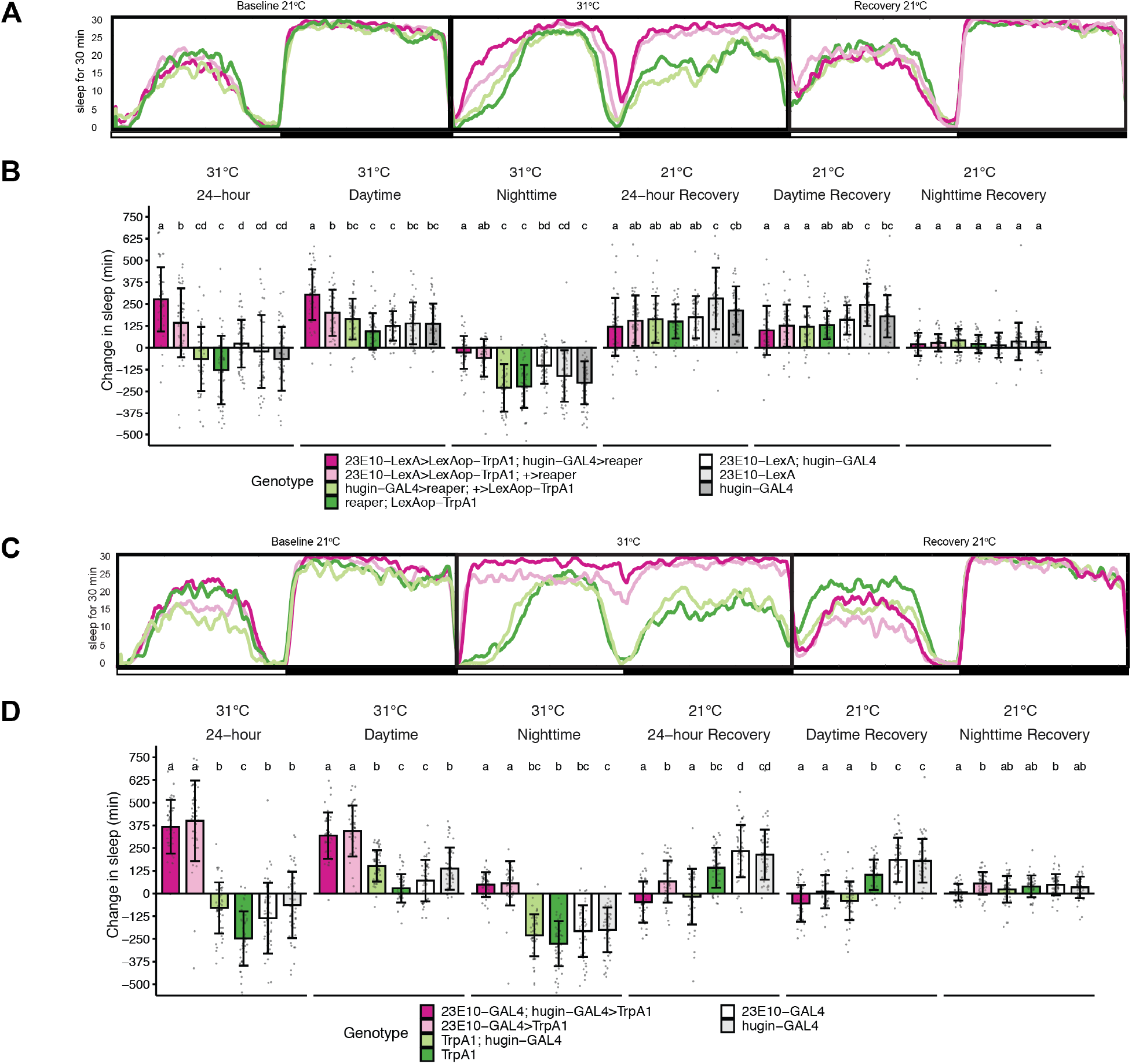
*hugin*+ neurons are effectors of *23E10*+ sleep-promoting dFB neurons. **(A)** Sleep profiles before, during, and after *23E10*+ dFB neuron activation with *TrpA1* in flies where *hugin*+ neurons were ablated using *reaper* (n=42-47). **(B)** Quantification of A. Left 3 graphs: Changes in sleep amount with temperature dependent (31°C) activation of *23E10*+ neurons, relative to sleep levels on the pre-activation day. Changes are shown for the 24h day and split into day and night sleep. B, right three graphs: Assay of recovery sleep following *TrpA1* activation of *23E10*+ dFB neurons in the absence of *hugin*+ neurons. The higher temperature, used to induce *TrpA1*, decreases sleep and results in higher sleep the following day. Recovery sleep is plotted as the difference in 24h sleep on the day after heat-induced sleep loss relative to sleep amount on the pre-activation day. **(C)** Sleep profiles from before, during, and after simultaneous activation of *23E10*+ dFB neurons and *hugin*+ neurons with *TrpA1* (n=41-48). **(D)** Quantification of C. Left 3 graphs: Changes in sleep amount with temperature dependent (31°C) activation of *23E10*+ neurons, relative to sleep levels on the pre-activation day. D, 3 right graphs: Recovery sleep following *TrpA1* activation of *23E10*+ dFB neurons and *hugin*+ neurons. Recovery sleep is plotted as the difference in 24h sleep on the day after heat-induced sleep loss relative to sleep amount on the pre-activation day. For all panels: Circles are individual fly data points, and summary statistics are displayed as mean + SD. Means compared with one-way ANOVA and Tukey’s test. For panel B,D: Means sharing the same letter are not significantly different from each other (P > 0.05, Tukey’s test).

In the thermogenetic sleep experiments, we also observed significant heat-induced sleep loss during the night in control animals (Figure 3A, third graph). This is consistent with the effect of high temperature, which reorganizes sleep such that daytime sleep increases and night-time sleep decreases. Indeed, effects of dFB activation on night-time sleep are often manifested as unchanged sleep levels relative to the decrease seen in controls. Heat-induced nighttime sleep loss also engages the homeostat resulting in sleep increase after flies are returned to the baseline temperature ^34^. To determine if *hugin*+ neurons affect heat-induced sleep loss or subsequent recovery, we maintained flies for a day at 31°C (high temperature), after which they were returned to 21°C (low temperature) to recover sleep. Recovery sleep was determined by comparing sleep on the recovery day with sleep on the baseline day, with both days at 21°C. Ablation of *hugin*+ neurons did not affect the amount of sleep loss at 31°C or the amount of recovery sleep at 21°C after heat-induced sleep loss (Figure 3A-B). Thermogenetic activation of *hugin*+ neurons also did not affect the amount of sleep loss at 31°C, when compared to the controls (Figure 3C-D). However, after return to 21°C, recovery sleep was decreased in flies where *hugin*+ cells were activated with *TrpA1*, compared to the control groups or flies with *23E10*>*TrpA1* activation (Figure 3C-D).

Despite having more sleep than controls during the high temperature, flies with activated *23E10*+ dFB neurons recovered additional sleep after the transition back to low temperature. However, flies subjected to simultaneous activation of *23E10*+ and *hugin*+ neurons showed decreased sleep recovery at 21°C, similar to that seen with *hugin*>*TrpA1* activation alone (Figure 3C-D). We hypothesize that homeostatic pressure triggered by heat-induced sleep loss inhibits the activity of *hugin*+ circadian neurons to promote recovery sleep. Activation of *hugin*+ neurons reduces this recovery.

To verify that heat-induced sleep loss affects the activity of *hugin*+ neurons, we used the CaLexA system to measure Ca^2+^ in *hugin*>*CaLexA-GFP;RFP* flies after a full day at 30°C. A control group, flies that were kept at a constant low temperature, was assayed at the same time of day as the heat-exposed group. Compared to the baseline night at 21°C, *hugin*>*CaLexA-GFP;RFP* flies (*n* = 16) lost an average 208.13 minutes of sleep (sd = 79.10) during the night at 30°C (Figure S4A). The heat-induced sleep loss was accompanied with a decreased CaLexA-GFP signal (normalized by RFP) in *hugin*+ neurons (Figure S4B), showing that sleep loss due to heat also inhibits the activity of *hugin*+ neurons.

### Mutations in the *hugin* locus affect sleep rebound

To determine if the ability of *hugin*+ neurons to affect the sleep homeostat is dependent on Hugin peptide signaling, we used CRISPR-CAS9 to produce mutant alleles of *hugin* that affect expression of one or both of its encoded neuropeptides, Hugin-γ and Pyrokinin 2 (PK2 ^35^. The *hugin^PK^* mutant contains a 1-base pair deletion that truncates the PK2 peptide, while the *hugin^ΔEx3^* mutant lacks the majority of exon 3 and is predicted to eliminate both neuropeptides. We backcrossed each CRISPR line for 5 generations to exclude potential off target mutations and then tested the flies for circadian and sleep behavior. The *hugin^PK2^* mutants showed weaker circadian rhythms, while the *hugin^ΔEx3^* flies showed a small lengthening of circadian period (Figure 4C); thus, period effects may arise from the Hugin peptide, and we suggest that the stronger rhythms in the null allele reflect some developmental compensation. Interestingly, both mutants exhibited decreases in baseline sleep (Figure 4A). Since our data support the idea that *hugin*+ neurons are effectors of the sleep homeostat, we measured rebound sleep in both mutants after mechanical deprivation of sleep for nine hours at the end of the night (ZT 15-24). We found that both mutants exhibited increased rebound sleep (Figure 4B) indicating that effects of *hugin*+ neurons on recovery sleep are mediated through peptides made by the *hugin* locus.

**Figure 4:**
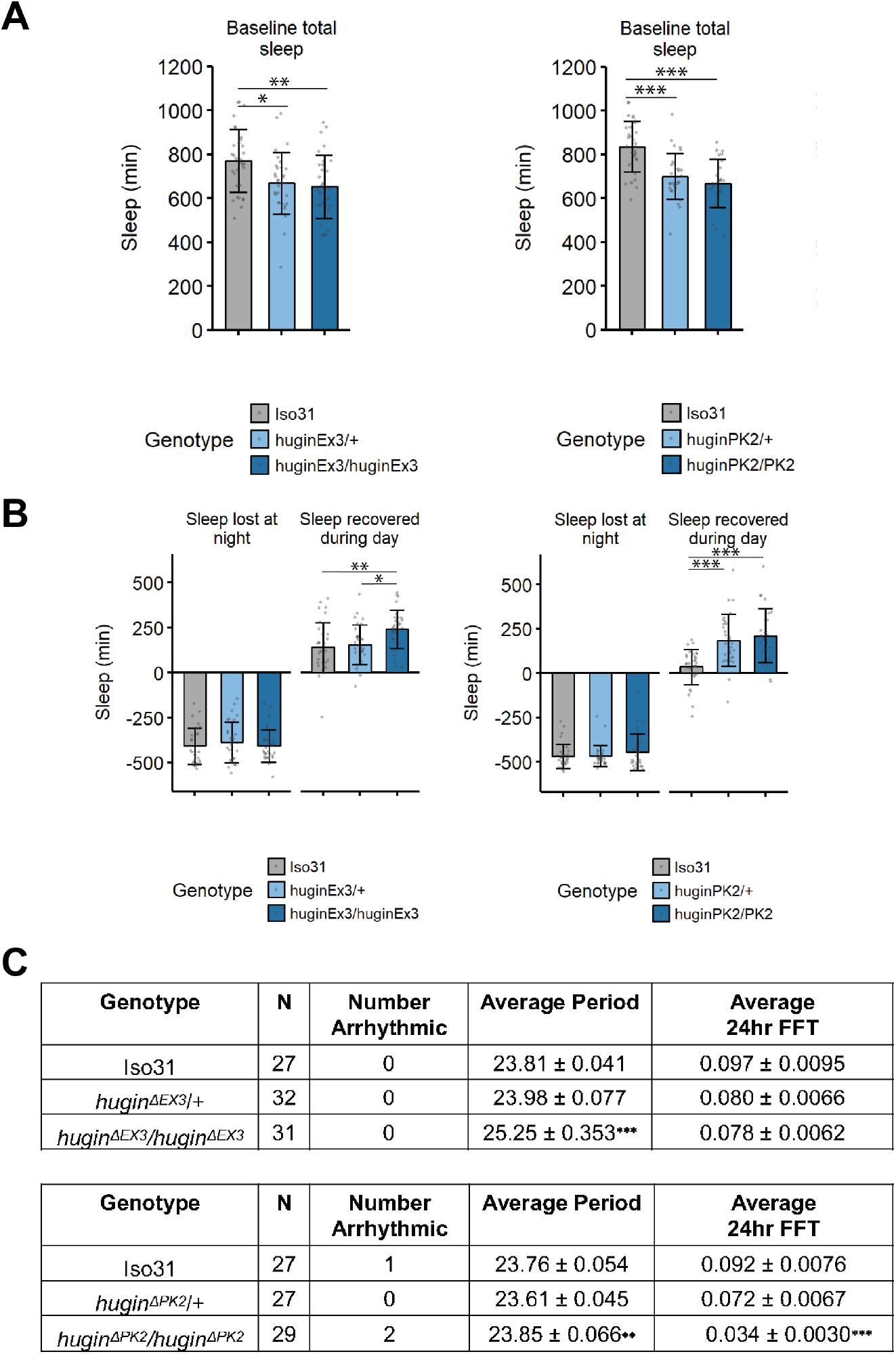
Hugin peptide signaling regulates baseline sleep and circadian rhythms. Hugin CRISPR mutants, *hugin^ΔEx3^* (left A,B) and *hugin^PK^*(right A,B), were assayed for baseline sleep **(A)** and response to mechanical sleep deprivation from ZT15-24 **(B)**. In B, sleep was assayed during and following deprivation. For Panels A-B: N=27-32 flies. Circles are individual fly data points, and summary statistics are displayed as mean + SD. Analysis was analyzed using one-way ANOVA and Tukey’s test. **P < 0.05, **P < 0.01, ***P < 0.001. **(C)** Hugin CRISPR mutations affect circadian rhythms. Top: A homozygous mutation for *hugin^ΔEx3^* increases average period length. *** P<0.0001 by Tukey’s post hoc test after a one-way ANOVA compared to Iso31 and *huginΔEX3/*+. Bottom: A homozygous mutation for the *hugin^ΔPK2^* decreases 24-hour rhythm strength. *** P<0.0001 by Tukey’s post hoc test after a one-way ANOVA compared to Iso31 and *huginΔPK2/*+. ^♦♦^ P<0.01 by Tukey’s post hoc test after a oneway ANOVA compared to *huginΔPK2/*+.

### *Pdf*+ clock neurons are targets of *hugin*+ neurons

We previously showed that *hugin*+ neurons project into the ventral nerve cord^7^. To identify additional targets of these neurons, we used *trans*-Tango, a pan-neuronal transsynaptic labeling system in which a tethered ligand at synapses activates tdTomato expression in postsynaptic partners^36^. Presynaptic neurons are simultaneously labeled with myr::GFP, a different fluorescent protein. We expressed the *trans*-Tango ligand in *hugin*+ neurons and observed *trans*-Tango-dependent signal in many brain regions, including the pars intercerebralis, mushroom body lobes, mushroom body calyx and pedunculus, SMP, subesophageal zone, and accessory medulla (Figure 5A top). The postsynaptic neurons, with cell bodies in the accessory medulla and projections to the superior medial protocerebrum, were reminiscent of small ventrolateral clock neurons (s-LNvs) (Figure 5A top right, magenta, arrowheads). S-LNvs secrete the peptide Pigment Dispersing Factor (PDF), prompting us to label for PDF peptide and confirm that a subset of the postsynaptic partners observed in *hugin*>*trans-Tango* flies is PDF-positive. *Pdf*+ neurons include the large ventrolateral neurons (l-LNv), which were also positive for the *trans*-Tango-dependent signal, but showed weaker signal than the s-LNvs (Figure 5A bottom), indicating that s-LNvs are primary targets of *hugin*+ neurons. To confirm the synaptic connection between *hugin*+ and *PDF*+ neurons, we used GRASP as above. Expression of synaptic nrx::spGFP1-10 in *hugin*+ neurons and spGFP11 in *PDF*+ neurons resulted in GFP reconstitution in the superior medial protocerebrum (SMP), superior lateral protocerebrum (SLP), and posterior lateral protocerebrum (PLP) (Figure 5B). Thus, two independent techniques identified a synaptic connection localized to the same part of the brain, indicating that *hugin*+ neurons indeed project to *Pdf*+ neurons.

**Figure 5:**
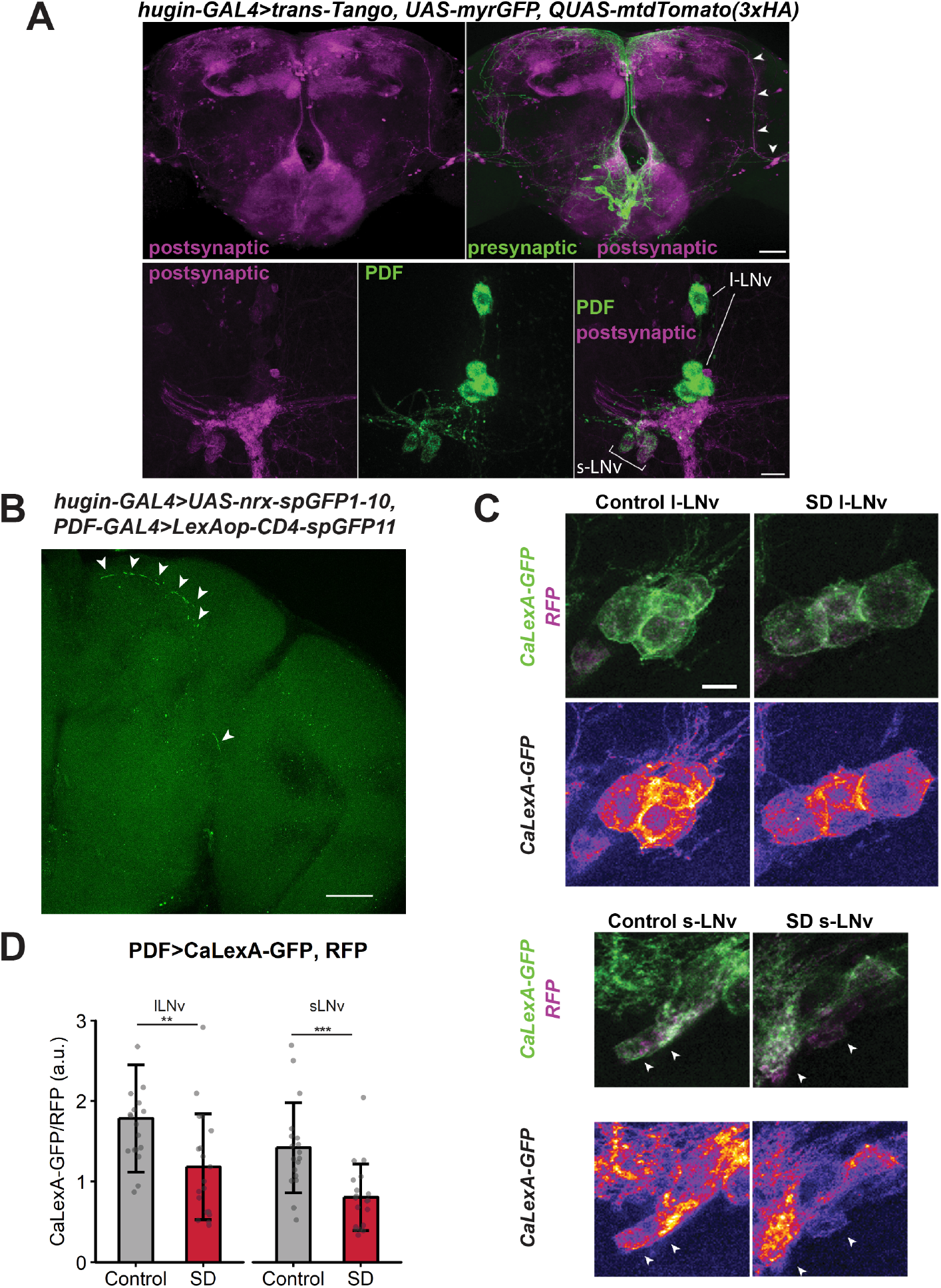
*hugin*+ neurons target PDF-expressing clock neurons which show decreased Ca^2+^ levels upon sleep deprivation. **(A)** Use of *trans*-Tango to map projections of *hugin*+ neurons (green). *trans*-Tango system reveals the synaptic partners (magenta) of *hugin*+ neurons in the brain. Panel A (top) image is a max intensity projection from the anterior side, and arrowheads indicates postsynaptic signal that resembles the projections of PDF+ neurons. A (bottom) Co-labeling of PDF peptide (green) and postsynaptic signal (magenta) in flies with *trans*-Tango ligand expressed in *hugin*+ neurons. PDF+ s-LNvs are postsynaptic to *hugin*+ neurons. s-LNv, small ventrolateral neurons, l-LNv, large ventrolateral neurons. Scale bars, A top: 50 μm; bottom: 15 μm. **(B)** GRASP to assay connectivity between *hugin*+ neurons and LNvs. Synaptic nrx::spGFP1-10 is expressed in presynaptic neurons and complementary spGFP11 expressed in putative postsynaptic neurons. GFP reconstitution occurs only if synaptic connectivity exists. Expression of nrx::spGFP1-10 in *hugin*+ neurons and spGFP11 in *PDF*+ neurons resulted in GFP reconstitution (white arrows) occurs in the superior medial protocerebrum (SMP), superior lateral protocerebrum (SLP), and posterior lateral protocerebrum (PLP). Scale bar 50 μm. **(C)** Representative images of l-LNvs or s-LNvs from a *Pdf*>*CaLexA-GFP; RFP* fly in Control or SD group. Top row shows merged images of GFP signal reporting Ca^2+^ levels with the CaLexA system and RFP normalizer signal. Bottom row shows “Fire” pseudocolor image of CaLexA-GFP signal (blue/purple=low intensity and yellow/white=high intensity). Scale bar, 10 μm applies for all images in this panel. **(D)** *Pdf*>*CaLexA-GFP; RFP* flies were subjected to no sleep deprivation (Control, gray) or 9-hr sleep deprivation (SD, red). Graph shows GFP levels normalized to RFP levels in *Pdf*+ large ventrolateral neurons (l-LNv) or small ventrolateral neurons (s-LNv) from Control (*n* = 18 flies) and SD (*n* = 19 flies) groups. **P = 0.00910, ***P = 0.000655 by Welch’s *t*-test.

Our data identify a circuit that links sleep homeostasis centers to circadian clock neurons (*23E10*+ dFB → *hugin*+ SEZ → *Pdf*+ s-LNvs), and suggest a potential mechanism for homeostatic components to regulate circadian clock outputs. To test whether the activity of *Pdf*+ neurons themselves is altered with sleep deprivation, we used CaLexA to measure Ca^2+^ level changes in *Pdf*+ neurons during sleep deprivation. With mechanical sleep deprivation, the CaLexA-GFP signal in both *Pdf*+ s-LNv and l-LNv cell bodies was lower in the sleep-deprived flies as compared to controls (Figure 5C,D). We next tested if the Ca^2+^ decrease in *Pdf*+ neurons after deprivation is dependent on Hugin peptide signaling. By comparing the activity of *Pdf*+ neurons in control and sleep deprivation conditions, we found that in s-LNvs, sleep deprivation resulted in a decrease in Ca^2+^ levels in the control *hugin^ΔEx3/+^* group, but not in *hugin^ΔEx3/ΔEx3^* mutants (Figure 6A,B). This result is of particular interest since the *trans*-Tango experiment suggests that s-LNvs are primary targets of *hugin*+ neurons. While sleep deprivation did not significantly affect Ca^2+^ levels in l-LNvs of *hugin* mutants or *hugin^ΔEx3/+^* controls, we suspect that the large area of the large neurons dilutes changes in calcium, so we cannot definitively say that only s-LNvs show consistent responses to sleep loss. Together these data indicate that sleep deprivation acts through Hugin peptides to suppress the activity of LNvs.

**Figure 6:**
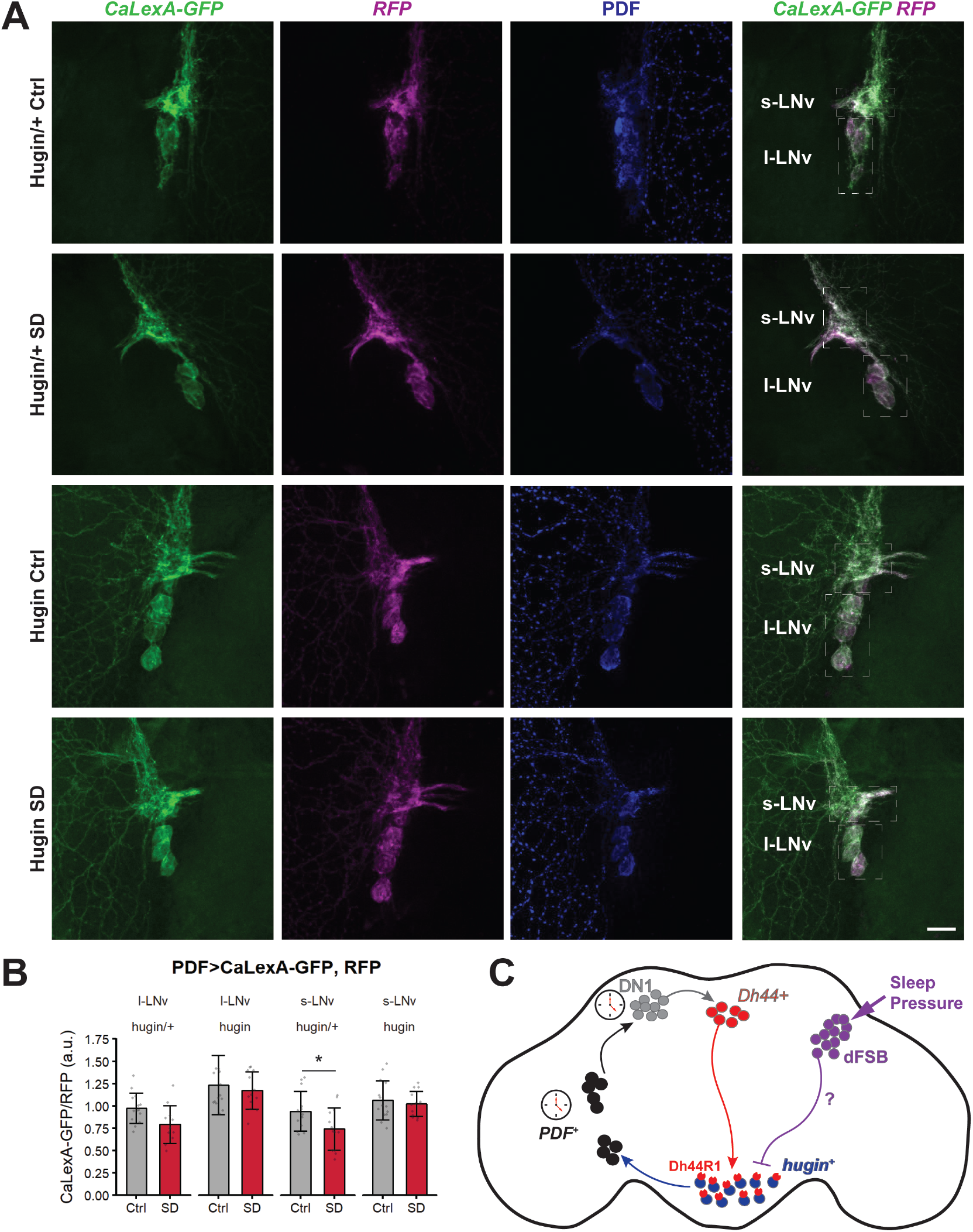
Hugin peptide mediates the decrease in Ca^2+^ levels of *Pdf*-expressing clock neurons after sleep deprivation. **(A)** Representative images of l-LNvs and s-LNvs from a *Pdf*>*CaLexA-GFP; RFP* flies in Control or SD group. The fourth column shows merged images of GFP signal reporting Ca^2+^ levels with the CaLexA system and RFP normalizer signal. Note the lower green signal in the *hugin^ΔEx3/+^* heterozygous fly subjected to SD (second row) compared to the *hugin^ΔEx3/+^* heterozygous control fly (top row). 25 μm scale bar applies for all images in this panel **(B)** Ca^2+^ levels in *Pdf*+ neurons were measured with *Pdf*>*CaLexA-GFP; RFP* reporter in *hugin^ΔEx3/+^* heterozygous or *hugin^ΔEx3/ΔEx3^* mutant flies that were subjected to no sleep deprivation (Control, gray) or 9-hr sleep deprivation (SD, red). Graph shows GFP levels normalized to RFP levels in *Pdf*+ l-LNvs or s-LNvs (*n* = 13 or 14 flies per group). l-LNv: Two-way ANOVA (genotype × condition) revealed a significant main effect of genotype (F(1, 47) = 23.29, P < 0.0001), a nonsignificant effect of condition (F(1, 47) = 3.68, P = 0.061), and a nonsignificant interaction between factors (F(1, 47) = 0.604, P = 0.441). s-LNv: Two-way ANOVA revealed a significant main effect of genotype (F(1, 49) = 12.47, P = 0.0009), a significant effect of condition (F(1, 49) = 4.36, P = 0.042), and a nonsignificant interaction between factors (F(1, 49) = 1.852, P=0.180). * P = 0.0385, significantly different by Sidak’s multiple comparison test. **(C)** Proposed model for regulation of a circadian output circuit by sleep homeostatic drive. Sleep drive triggered by prolonged wakefulness suppresses locomotor activity by decreasing firing of *hugin*+ neurons (blue) and, through them, also firing of *PDF*+ clock neurons (black). We suggest that effects of the sleep homeostat on hugin+ neurons are mediated by projections of *23E10*+ dFB neurons (purple).

## Discussion

The circadian clock and homeostat both regulate sleep, but it is not clear how the two processes functionally interact. We show here that circadian and sleep circuits intersect at circadian output neurons that secrete the Hugin peptide. Homeostatic sleep drive signals through *hugin*+ neurons to suppress circadian outputs, thereby allowing for sleep at times when the circadian system typically promotes wake. We also find that *hugin*+ circadian output neurons feedback to s-LNvs, the central clock neurons. Thus, the sleep homeostat influences outputs of the circadian clock by modulating the activity of circadian output neurons and clock neurons (Figure 6C).

Previously, we showed that a circuit from s-LNvs → DN1 → *Dh44*+ neurons → *hugin*+ neurons controls rhythms of locomotor activity. *hugin*+ neurons promote locomotion in the morning and evening, but especially in the evening during the day-to-night transition ^6,7^. Ablation of these neurons does not affect baseline sleep, suggesting that their primary function is in the modulation of locomotor activity; however, they are relevant for homeostatic responses to sleep loss. As for *hugin*+ neurons, the major function of the dFB *23E10*+ neurons is in the context of sleep loss. Given that *23E10*+ neurons project to *hugin*+ neurons and are activated by sleep loss while *hugin*+ neurons show decreased activity ^24^, we suggest that *23E10*+ neurons inhibit *hugin*+ neurons. This would also explain why ablation of *hugin*+ neurons enhances sleeppromoting effects of *23E10*+ activation. We were unable to reliably detect inhibition of *hugin*+ neurons by *23E10*+ neurons through optogenetic or other stimulus-response experiments (data not shown), but this could be due to heterogeneity in the *hugin*+ population. Regardless of whether the dFB is their major source of homeostatic input, it is clear that *hugin*+ neurons affect sleep homeostasis. Activating *hugin*+ neurons during a period of heat-induced sleep loss decreases recovery sleep, and while it does not affect rebound following mechanical deprivation, this is likely because mechanical deprivation recruits multiple pathways ^21^. We note also that in the mechanical sleep deprivation experiment, *hugin*+ neurons were activated prior to deprivation, while heat-induced sleep loss was concurrent with their activation; it is possible that prior activation resets the homeostatic system to a different set point of neural activity. Importantly, we also report increased recovery sleep in *hugin* mutants following deprivation, indicating that the homeostatic sleep-suppressing role of *hugin*+ neurons is mediated by the Hugin peptide. Our data suggest that during sleep deprivation, the homeostat not only generates sleep drive but also actively disengages activity-promoting circuits.

The circadian clock can regulate sleep by cell-intrinsically controlling the neuronal activity of clock neurons, such as the LNvs. The wake-promoting effect of the LNvs is lightdependent and largely comes from the l-LNv subset ^37–39^. While s-LNvs alone are not sufficient to promote wake, downregulation of PDF receptor in s-LNvs increases sleep, suggesting PDF signaling to s-LNvs modulates wake-promoting effects of l-LNvs ^38,39^. In addition, the downregulation of short Neuropeptide F signaling between s-LNvs and l-LNvs decreases nighttime sleep ^40^. Notably, both s-LNvs and l-LNvs show more depolarized resting membrane potentials during the day than during the night, supporting the idea that LNvs are more active during times of increased arousal ^41,42^. A recent study demonstrated that another subset of clock neurons, LPN^AstA^ neurons, projects to the dFB to promote sleep and could serve as a mechanism of circadian control of the sleep homeostat ^20^.

Our work suggests that sleep homeostasis also influences the neuronal activity of the LNvs. An increase in sleep drive caused by social enrichment was previously associated with an increased number of synapses in the LNv projections into the medulla, a brain region that processes visual information from the eyes ^43^. Here, we report Ca^2+^ levels in LNvs decrease with sleep deprivation, which we hypothesize dampens the wake-promoting effects of LNvs to allow for recovery sleep. It is possible that decreased Ca^2+^ levels in LNvs with sleep deprivation precede synaptic downscaling that is thought to occur with recovery sleep ^43^. *hugin* mutants do not exhibit a decrease in s-LNv *Ca2*+ levels with sleep deprivation, which suggests that Hugin peptide signaling is involved in the suppression of s-LNv activity with sleep deprivation. Although we demonstrate projections from the dFB to *hugin*+ neurons, other sleep homeostat pathways may also modulate *hugin*+ neurons and/or LNvs. Notably, GABA and myoinhibitory peptide signal to LNvs to regulate sleep, although the source of these neuromodulators is not known yet ^39,44,45^.

The mammalian orthologue of Hugin, Neuromedin U (NMU) ^35^, is also implicated in sleep regulation. While NMU injection in rats does not change total sleep time, it changes sleep architecture ^46^. NMU overexpression decreases sleep in zebrafish ^47^, which is consistent with our findings that *hugin*+ neurons promote wake. Chiu et. al did not report alterations in baseline sleep levels in their NMU knockout fish, but we find that *hugin* CRISPR mutants exhibit decreased baseline sleep. We do not have an explanation for the reduced baseline sleep, but note that following deprivation, *hugin* mutants show increased rebound, supporting the idea that Hugin opposes increases in sleep following deprivation. Interestingly, NMU overexpression in zebrafish increases the duration of stimulus-evoked arousal during sleep. To the extent that increased arousal reflects decreased sleep drive, this indicates a conserved role for Hugin/NMU in regulating sleep.

In *Drosophila*, influences of the sleep homeostat on the circadian system have previously not been demonstrated. In rodents, sleep deprivation dampens electrical activity in the suprachiasmatic nucleus (SCN) for up to 7 hours and reduces the ability of the circadian clock to phase shift by light ^1,48–50^. We find a similar effect in flies, where neuronal activity is depressed in LNv central clock neurons and remained depressed even 5 hours after the deprivation ended (data not shown). In the rodent model, the mechanism mediating the reduced SCN activity is not clear, but may involve serotonin signaling from the raphe dorsalis ^51^. We show here that effects of sleep deprivation on LNvs are mediated by Hugin signaling from output neurons that connect to a major sleep homeostatic locus. Importantly, sleep deprivation does not appear to shift rest:activity patterns in flies ^52^ and nor does it affect clock gene expression in the rodent SCN ^53^, suggesting that timekeeping by the clock is relatively unperturbed. Therefore, sleep homeostasis appears to primarily influence clock outputs.

## Methods

### Drosophila melanogaster

Flies were maintained on cornmeal-molasses medium. For thermogenetic and *trans*-Tango experiments, flies were raised at 18°C, and all other flies were maintained at 25°C. *w^1118^* iso31 strain was used as the wild type strain. For sleep behavior experiments, transgenic lines were backcrossed into the iso31 genetic background. For controls, UAS and GAL4 fly lines were tested as heterozygotes after crossing to iso31. The following flies were from the Bloomington Drosophila Stock Center: *23E10-GAL4* (#49032) ^54^, *23E10-LexA* (#52693) ^55^, *Hugin-GAL4* (#58769) ^56^, *Hugin-LexA* (#52715), *Dh44-GAL4* (#39347), *UAS-CD8::RFP* (#32219), *LexAop-Rab3::GFP* (#52239) ^57^, *LexAop-6xmCherry-HA* (#52271), *UAS-nSyb::GFP1-10, LexAop-CD4::GFP11* (#64314), *UAS-reaper* (#5773) ^58^. *LexAop-CD4-spGFP11; UAS-nrx-spGFP1-10* was a gift from N. Shah ^59^. *Trans*-Tango fly was a gift from G. Barnea ^36^. CaLexA fly was a gift from J.W. Wang ^25^. *PDF-GAL4* was a gift from J. Hall ^60^. *UAS-TrpA1* was a gift from L.C. Griffith ^18^. *UAS-shibire^ts^* (*20XUAS-IVS-Shibire[ts1]-p10-INS*) and *LexAop-TrpA1* (chromosome 2) were gifts from G. Rubin ^61^. *LexAop-TrpA1* (chromosome 3) was a gift from S. Waddell ^62^. The *LexAop-TrpA1; LexAop-TrpA1* fly line was created by H. Toda (Sehgal Lab).

### Generation of *hugin* mutants

*hugin^ΔEx3^* and *hugin^PK2^* mutants were generated with the CRISPR-CAS9 system. The guide RNA sequences for generating *hugin^ΔEx3^* were 5’ GGGAGCCCGCTTATCGCGTG 3’ and 5’ GGAGGACGGAGGACGAGCCC 3’, and the guide RNAs for *hugin^PK2^* were 5’ GTGCCGTTCAAGCCACGCCT 3’ and 5’ GGCAAACGTGCTCAAGTGTG 3’. Guide RNAs were cloned into the pCFD4 plasmid ^63^, and the plasmids encoding the guide RNAs were injected into *vasa-Cas9* flies (BDSC # 51323) ^64^ at Rainbow Transgenic Flies, Inc (Camarillo, CA). Mutations in the F1 generation were identified with PCR screening and confirmed with Sanger sequencing. *hugin^ΔEx3^* is a 260-bp deletion (dm6/chr3R:12,528,601-12,528,860), and *hugin^PK2^* is a 1-bp deletion (dm6/chr3R: 12,528,750). The mutant alleles were backcrossed for five generations into iso31 background.

### Immunohistochemistry

For polarity labeling, CaLexA experiments, and reaper confirmation ~7 d old females raised at 25°C were used. For *trans*-Tango experiments, ~15-20 d old females raised at 18°C were used, as previously described ^36^. All fly brains were dissected in phosphate-buffered saline with 0.1% Triton-X (PBST) and fixed with 4% formaldehyde in PBS for 20 min at room temperature. Brains were rinsed 3 × 10 min with PBST, blocked in 5% Normal Goat Serum in PBST (NGST) for 60 min, and incubated in primary antibody diluted in NGST for >16 h at 4°C. Brains were rinsed 3 × 10 min in PBST, incubated 2 h in secondary antibody diluted in NGST, rinsed 3 × 10 min in PBST, and mounted with Vectashield media (Vector Laboratories Inc.). For the reaper confirmation experiment both primary and secondary antibody incubation was for two nights at 4°C. Primary antibodies used were: rabbit anti-GFP at 2μg/mL (Thermo Fisher Scientific Inc. A-11122), rat anti-RFP at 1μg/mL (ChromoTek 5F8), mouse anti-BRP at 1:100 (Developmental Studies Hybridoma Bank nc82), rat anti-HA at 1μg/mL (Roche clone 3F10), and mouse anti-PDF at 0.3μg/mL (Developmental Studies Hybridoma Bank c7-c). Secondary antibodies were from Thermo Fisher Scientific Inc. and used at 1:1000: Alexa Fluor 488 goat anti-rabbit, Alexa Fluor 555 goat anti-rat, Alexa Fluor 647 goat anti-rat, Alexa Fluor 647 goat anti-mouse.

### GRASP

nSyb-GRASP flies were dissected in extracellular saline (103 mM NaCl, 3 mM KCl, 1 mM NaH_2_PO_4_, 4 mM MgCl_2_, 10 mM D-(+)-trehalose dehydrate, 10 mM D-(+)-glucose, 5 mM N-tris(hydroxymethyl) methyl-2-aminoethane sulfonic acid, 26 mM NaHCO_3_, pH 7.4). Dissected brains were exposed to a high concentration of KCl to increase GRASP signal, as previously described ^32^. Dissected brains were incubated in 1 ml 70 mM KCl in saline three times (~5 s per KCl incubation), alternating with 1 ml saline (~5 s per wash), and then transferred to 1 mL saline to incubate for 10 minutes. Brains were fixed with 4% formaldehyde in PBS for 20 minutes at room temperature, rinsed 3 × 10 min in PBST, and mounted with Vectashield media. nrx-GRASP brains were processed using the same protocol except without the KCl washes. Endogenous GRASP signal without antibody labeling was imaged.

### Confocal Microscopy

Eight-bit images were acquired using a Leica TCS SP5 laser scanning confocal microscope with a 40x/1.3 NA or 20x/0.7 NA objective and a 1-μm or 2-μm z-step size. Maximum intensity z-projection and sum intensity z-projection (reaper confirmation only) images were generated in Fiji, a distribution of ImageJ software ^65^.

### Sleep Behavior Assay

Individual ~7 d old female flies were loaded into glass tubes containing 5% sucrose and 2% agar. Locomotor activity was monitored with the Drosophila Activity Monitoring system (DAMS) (Trikinetics, Waltham, MA). Flies were monitored for sleep in a 12 h:12 h (12:12) light:dark cycle at 25°C for CaLexA experiments or at 21°C for thermogenetic experiments. Incubator temperature shifts occurred at lights-on, Zeitgeber time (ZT) 0. For mechanical sleep deprivation experiments, flies were loaded into the DAMS and sleep deprived during the night by shaking on an adapted vortex for 2 s randomly within every 20 s interval. Sleep was defined as 5 consecutive min of inactivity. Sleep analysis was performed with PySolo software ^66^. Data from flies that survived the duration of the experiments were pooled and analyzed. Behavioral data were analyzed with one-way analysis of variance (ANOVA) with Tukey’s test as the post hoc test to compare means between groups. Differences between groups were considered significant if P < 0.05 by the post hoc test.

### Circadian Rest:Activity Rhythms Behavior Assay

Rest:activity rhythm assays were performed and analyzed as described in King et. al. 2017. Flies were entrained to a 12 hr light: 12 hr dark (LD) cycle from birth and put in total darkness for analysis at ~7 d old. Data from days 3-9 for flies that survived the duration of the experiment were analyzed. Period and 24hr FFT were analyzed by one-way ANOVA with Tukey’s test in Prism 8 for Mac Os X.

### CaLexA Analysis

Fluorescence intensity measurement was performed in Fiji. Regions of interest (ROIs) were manually drawn to encompass individual RFP-positive cell bodies, and mean pixel intensities of RFP and GFP were measured from the ROI. For each cell, the CaLexA-GFP/RFP signal (arbitrary unit, a.u.) was calculated as a ratio between the mean pixel intensities of GFP and RFP. For each brain, the average CaLexA-GFP/RFP signal of a cell is a sample point. Welch’s *t*-test was used to compare differences in CaLexA-GFP/RFP signal between sleep-deprived and control groups. The two-way ANOVA was used to compare differences between groups that were split into genotype and conditions, and the Sidak’s multiple comparison post-hoc test was used for pairwise comparisons.

### Statistical Analysis

The statistical details of experiments can be found in figure legends. All statistical tests were performed in R (version 3.3.1). Graphs were generated in R using ggplot2 package, except for sleep profiles, which were generated in Pysolo.

## Acknowledgements

We thank Drs. Gilad Barnea, Leslie Griffith, Gerald Rubin, Jeffery Hall, Nirao Shah, Scott Waddell, and Jing Wang for generously providing fly lines. Also, we used stocks from the Bloomington Drosophila Stock Center (NIH P40OD018537). We thank Zhifeng Yue, Kiet Luu, and Juliana T. Choi for their assistance with experiments. The work was supported by grant NIH R37NS048471 (to A.S.). J.E.S. was supported by a training grant in Neuroscience (NIH T32-NS105607), an NIH Diversity Supplement (NIH NS48471), and by a grant to the University of Pennsylvania from the Howard Hughes Medical Institute through the James H. Gilliam Fellowship for Advanced Study program. A.N.K. was supported in part by a training grant in Genetics (NIH T32GM008216) and an NRSA fellowship (NIH F31NS100395). C.T.H. was supported by a training grant in Age Related Neurodegenerative Diseases (NIH T32AG00255).

## Author Contributions

Conceptualization, A.N.K. and A.S.; Methodology, J.E.S, A.N.K., C.T.H., A.F.B., and A.S.; Formal Analysis, J.E.S. and A.N.K.; Investigation, J.E.S., A.N.K., and C.T.H.; Resources, A.F.B; Writing – Original Draft, J.E.S., A.N.K., and A.S.; Writing – Review & Editing, J.E.S., A.N.K., C.T.H., A.F.B. and A.S.; Visualization, J.E.S., A.N.K., and C.T.H.; Supervision, A.S.

## References

1. Borbély, A. A., Daan, S., Wirz-Justice, A. & Deboer, T. The two-process model of sleep regulation: A reappraisal. J. Sleep Res. 25, 131–143 (2016).

2. Dubowy, C. & Sehgal, A. Circadian Rhythms and Sleep in Drosophila melanogaster. Genetics 205, 1373–1397 (2017).

3. Schlichting, M., Díaz, M. M., Xin, J. & Rosbash, M. Neuron-specific knockouts indicate the importance of network communication to drosophila rhythmicity. Elife (2019). doi:10.7554/eLife.48301

4. Delventhal, R. et al. Dissection of central clock function in drosophila through cell-specific CRISPR-mediated clock gene disruption. Elife (2019). doi:10.7554/eLife.48308

5. King, A. N. & Sehgal, A. Molecular and circuit mechanisms mediating circadian clock output in the Drosophila brain. European Journal of Neuroscience (2020). doi:10.1111/ejn.14092

6. Cavanaugh, D. J. et al. Identification of a circadian output circuit for rest:activity rhythms in Drosophila. Cell 157, 689–701 (2014).

7. King, A. N. et al. A Peptidergic Circuit Links the Circadian Clock to Locomotor Activity. Curr. Biol. (2017). doi:10.1016/j.cub.2017.05.089

8. Bai, L. et al. A Conserved Circadian Function for the Neurofibromatosis 1 Gene. Cell Rep. (2018). doi:10.1016/j.celrep.2018.03.014

9. Cavey, M., Collins, B., Bertet, C. & Blau, J. Circadian rhythms in neuronal activity propagate through output circuits. Nat. Neurosci. 19, 1–11 (2016).

10. Joiner, W. J., Crocker, A., White, B. H. & Sehgal, A. Sleep in Drosophila is regulated by adult mushroom bodies. Nature 441, 757–60 (2006).

11. Pitman, J. L., McGill, J. J., Keegan, K. P. & Allada, R. A dynamic role for the mushroom bodies in promoting sleep in Drosophila. Nature 441, 753–6 (2006).

12. Sitaraman, D. et al. Propagation of Homeostatic Sleep Signals by Segregated Synaptic Microcircuits of the Drosophila Mushroom Body. Curr. Biol. 25, 2915–2927 (2015).

13. Donlea, J. M. Neuronal and molecular mechanisms of sleep homeostasis. Curr. Opin. Insect Sci. 24, 51–57 (2017).

14. Donlea, J. M., Thimgan, M. S., Suzuki, Y., Gottschalk, L. & Shaw, P. J. Inducing sleep by remote control facilitates memory consolidation in Drosophila. Science 332, 1571–6 (2011).

15. Ueno, T. et al. Identification of a dopamine pathway that regulates sleep and arousal in Drosophila. Nat. Neurosci. 15, 1516–23 (2012).

16. Qian, Y. et al. Sleep homeostasis regulated by 5HT2b receptor in a small subset of neurons in the dorsal fan-shaped body of drosophila. Elife 6, (2017).

17. Liu, S., Liu, Q., Tabuchi, M. & Wu, M. N. Sleep Drive Is Encoded by Neural Plastic Changes in a Dedicated Circuit. Cell 165, 1347–1360 (2016).

18. Pulver, S. R., Pashkovski, S. L., Hornstein, N. J., Garrity, P. A. & Griffith, L. C. Temporal dynamics of neuronal activation by Channelrhodopsin-2 and TRPA1 determine behavioral output in Drosophila larvae. J. Neurophysiol. 101, 3075–88 (2009).

19. Kitamoto, T. Conditional modification of behavior in Drosophila by targeted expression of a temperature-sensitive shibire allele in defined neurons. J. Neurobiol. 47, 81–92 (2001).

20. Ni, J. D. et al. Differential regulation of the Drosophila sleep homeostat by circadian and arousal inputs. Elife (2019). doi:10.7554/eLife.40487

21. Dubowy, C. et al. Genetic Dissociation of Daily Sleep and Sleep Following Thermogenetic Sleep Deprivation in Drosophila. Sleep 39, 1083–95 (2016).

22. Bushey, D., Tononi, G. & Cirelli, C. Sleep- and wake-dependent changes in neuronal activity and reactivity demonstrated in fly neurons using in vivo calcium imaging. Proc. Natl. Acad. Sci. U. S. A. 112, 4785–90 (2015).

23. Yap, M. H. W. et al. Oscillatory brain activity in spontaneous and induced sleep stages in flies. Nat. Commun. 8, 1815 (2017).

24. Donlea, J. M., Pimentel, D. & Miesenbock, G. Neuronal machinery of sleep homeostasis in Drosophila. Neuron 81, 860–872 (2014).

25. Masuyama, K., Zhang, Y., Rao, Y. & Wang, J. W. Mapping neural circuits with activity-dependent nuclear import of a transcription factor. J. Neurogenet. 26, 89–102 (2012).

26. Pimentel, D. et al. Operation of a homeostatic sleep switch. Nature 536, 333–337 (2016).

27. Donlea, J. M. et al. Recurrent Circuitry for Balancing Sleep Need and Sleep. Neuron 1–12 (2018). doi:10.1016/j.neuron.2017.12.016

28. Li, W. et al. Morphological characterization of single fan-shaped body neurons in Drosophila melanogaster. Cell Tissue Res. 336, 509–519 (2009).

29. Cavanaugh, D. J., Vigderman, A. S., Dean, T., Garbe, D. S. & Sehgal, A. The Drosophila Circadian Clock Gates Sleep through Time-of-Day Dependent Modulation of Sleep-Promoting Neurons. Sleep 39, 345–56 (2016).

30. Schmid, A. et al. Activity-dependent site-specific changes of glutamate receptor composition in vivo. Nat. Neurosci. 11, 659–66 (2008).

31. Fouquet, W. et al. Maturation of active zone assembly by Drosophila Bruchpilot. J. Cell Biol. 186, 129–45 (2009).

32. Macpherson, L. J. et al. Dynamic labelling of neural connections in multiple colours by trans-synaptic fluorescence complementation. Nat. Commun. 6, 10024 (2015).

33. Feinberg, E. H. et al. GFP Reconstitution Across Synaptic Partners (GRASP) defines cell contacts and synapses in living nervous systems. Neuron 57, 353–63 (2008).

34. Parisky, K. M., Agosto Rivera, J. L., Donelson, N. C., Kotecha, S. & Griffith, L. C. Reorganization of Sleep by Temperature in Drosophila Requires Light, the Homeostat, and the Circadian Clock. Curr. Biol. 26, 882–92 (2016).

35. Melcher, C., Bader, R., Walther, S., Simakov, O. & Pankratz, M. J. Neuromedin U and its putative Drosophila homolog hugin. PLoS Biol. 4, e68 (2006).

36. Talay, M. et al. Transsynaptic Mapping of Second-Order Taste Neurons in Flies by trans-Tango. Neuron 96, 783–795.e4 (2017).

37. Sheeba, V. et al. Large ventral lateral neurons modulate arousal and sleep in Drosophila. Curr. Biol. 18, 1537–45 (2008).

38. Shang, Y., Griffith, L. C. & Rosbash, M. Light-arousal and circadian photoreception circuits intersect at the large PDF cells of the Drosophila brain. Proc. Natl. Acad. Sci. U. S. A. 105, 19587–94 (2008).

39. Parisky, K. M. et al. PDF cells are a GABA-responsive wake-promoting component of the Drosophila sleep circuit. Neuron 60, 672–82 (2008).

40. Shang, Y. et al. Short neuropeptide F is a sleep-promoting inhibitory modulator. Neuron 80, 171–83 (2013).

41. Cao, G. & Nitabach, M. N. Circadian control of membrane excitability in Drosophila melanogaster lateral ventral clock neurons. J. Neurosci. 28, 6493–501 (2008).

42. Sheeba, V., Gu, H., Sharma, V. K., O’Dowd, D. K. & Holmes, T. C. Circadian- and light-dependent regulation of resting membrane potential and spontaneous action potential firing of Drosophila circadian pacemaker neurons. J. Neurophysiol. 99, 976–88 (2008).

43. Donlea, J. M., Ramanan, N. & Shaw, P. J. Use-dependent plasticity in clock neurons regulates sleep need in Drosophila. Science 324, 105–8 (2009).

44. Chung, B. Y., Kilman, V. L., Keath, J. R., Pitman, J. L. & Allada, R. The GABA(A) receptor RDL acts in peptidergic PDF neurons to promote sleep in Drosophila. Curr. Biol. 19, 386–90 (2009).

45. Oh, Y. et al. A Homeostatic Sleep-Stabilizing Pathway in Drosophila Composed of the Sex Peptide Receptor and Its Ligand, the Myoinhibitory Peptide. PLoS Biol. 12, e1001974 (2014).

46. Ahnaou, A. & Drinkenburg, W. H. I. M. Neuromedin U2 receptor signaling mediates alteration of sleep-wake architecture in rats. Neuropeptides (2011). doi:10.1016/j.npep.2011.01.004

47. Chiu, C. N. et al. A Zebrafish Genetic Screen Identifies Neuromedin U as a Regulator of Sleep/Wake States. Neuron (2016). doi:10.1016/j.neuron.2016.01.007

48. Mistlberger, R. E., Landry, G. J. & Marchant, E. G. Sleep deprivation can attenuate light-induced phase shifts of circadian rhythms in hamsters. Neurosci. Lett. 238, 5–8 (1997).

49. Challet, E., Turek, F. W., Laute, M.-A. & Van Reeth, O. Sleep deprivation decreases phase-shift responses of circadian rhythms to light in the mouse: role of serotonergic and metabolic signals. Brain Res. 909, 81–91 (2001).

50. Deboer, T., Détári, L. & Meijer, J. H. Long term effects of sleep deprivation on the mammalian circadian pacemaker. Sleep 30, 257–62 (2007).

51. Deboer, T. Sleep homeostasis and the circadian clock: Do the circadian pacemaker and the sleep homeostat influence each other’s functioning? Neurobiol. Sleep Circadian Rhythm. (2018). doi:10.1016/j.nbscr.2018.02.003

52. Hendricks, J. C. et al. Rest in Drosophila is a sleep-like state. Neuron 25, 129–38 (2000).

53. Curie, T., Maret, S., Emmenegger, Y. & Franken, P. In Vivo Imaging of the Central and Peripheral Effects of Sleep Deprivation and Suprachiasmatic Nuclei Lesion on PERIOD-2 Protein in Mice. Sleep 38, 1381–94 (2015).

54. Jenett, A. et al. A GAL4-driver line resource for Drosophila neurobiology. Cell Rep. 2, 991–1001 (2012).

55. Pfeiffer, B. D. et al. Refinement of tools for targeted gene expression in Drosophila. Genetics 186, 735–55 (2010).

56. Melcher, C. & Pankratz, M. J. Candidate gustatory interneurons modulating feeding behavior in the Drosophila brain. PLoS Biol. 3, e305 (2005).

57. Shearin, H. K., Dvarishkis, A. R., Kozeluh, C. D. & Stowers, R. S. Expansion of the gateway multisite recombination cloning toolkit. PLoS One 8, e77724 (2013).

58. White, K., Tahaoglu, E. & Steller, H. Cell Killing by the Drosophila Gene reaper. Science (80-.). 271, 805–807 (1996).

59. Fan, P. et al. Genetic and Neural Mechanisms that Inhibit Drosophila from Mating with Other Species. Cell 154, 89–102 (2013).

60. Park, J. H. Differential regulation of circadian pacemaker output by separate clock genes in Drosophila. Proc. Natl. Acad. Sci. (2000). doi:10.1073/pnas.070036197

61. Pfeiffer, B. D., Truman, J. W. & Rubin, G. M. Using translational enhancers to increase transgene expression in Drosophila. Proc. Natl. Acad. Sci. U. S. A. 109, 6626–31 (2012).

62. Burke, C. J. et al. Layered reward signalling through octopamine and dopamine in Drosophila. Nature 492, 433–7 (2012).

63. Port, F., Chen, H.-M., Lee, T. & Bullock, S. L. Optimized CRISPR/Cas tools for efficient germline and somatic genome engineering in Drosophila. Proc. Natl. Acad. Sci. U. S. A. 111, E2967–76 (2014).

64. Gratz, S. J. et al. Highly specific and efficient CRISPR/Cas9-catalyzed homology-directed repair in Drosophila. Genetics 196, 961–71 (2014).

65. Schindelin, J. et al. Fiji: an open-source platform for biological-image analysis. Nat. Methods 9, 676–82 (2012).

66. Gilestro, G. F. & Cirelli, C. pySolo: a complete suite for sleep analysis in Drosophila. Bioinformatics 25, 1466–7 (2009).

