## Supplemental Figures for "*Hugin*^+^ neurons link the sleep homeostat to circadian clock neurons"

### Supplemental Information

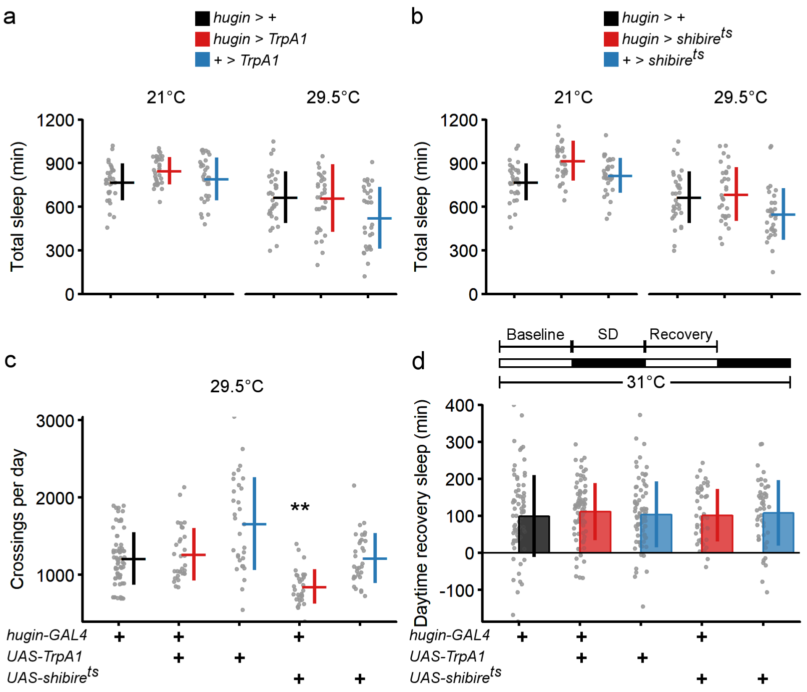


**Figure S1: Disrupting activity of *hugin*+ neurons does not alter normal sleep amount or recovery after mechanical sleep deprivation.**

**(A)** Thermogenetic activation of *hugin*+ neurons does not alter total amount of sleep. **(B)** Thermogenetic inhibition of synaptic transmission from *hugin*+ neurons does not alter total amount of sleep. Data for the *hugin*>+ control are the same for panels A and B as both manipulations involve testing at 29.5°C. **(C)** Blocking synaptic transmission in *hugin*+ neurons reduces the number of beaming crossing, a measure of locomotor activity. **(D)** Thermogenetic activation or inhibition of *hugin*+ neurons does not alter sleep recovery after mechanical sleep deprivation. Experimental setup: Flies were kept at 31°C for the entire duration of experiment. Flies were sleep deprived (SD) by mechanical shaking during the nighttime on day 1 and allowed to recover during the daytime on day 2. Change in day sleep was based upon sleep during 12 hours of the Recovery day relative to 12 hours of the Baseline day. $^{**}P < 0.01$ by

Tukey's test after one-way ANOVA. Circles are individual fly data points, and summary statistics displayed as mean + SD. N=32 for all groups.

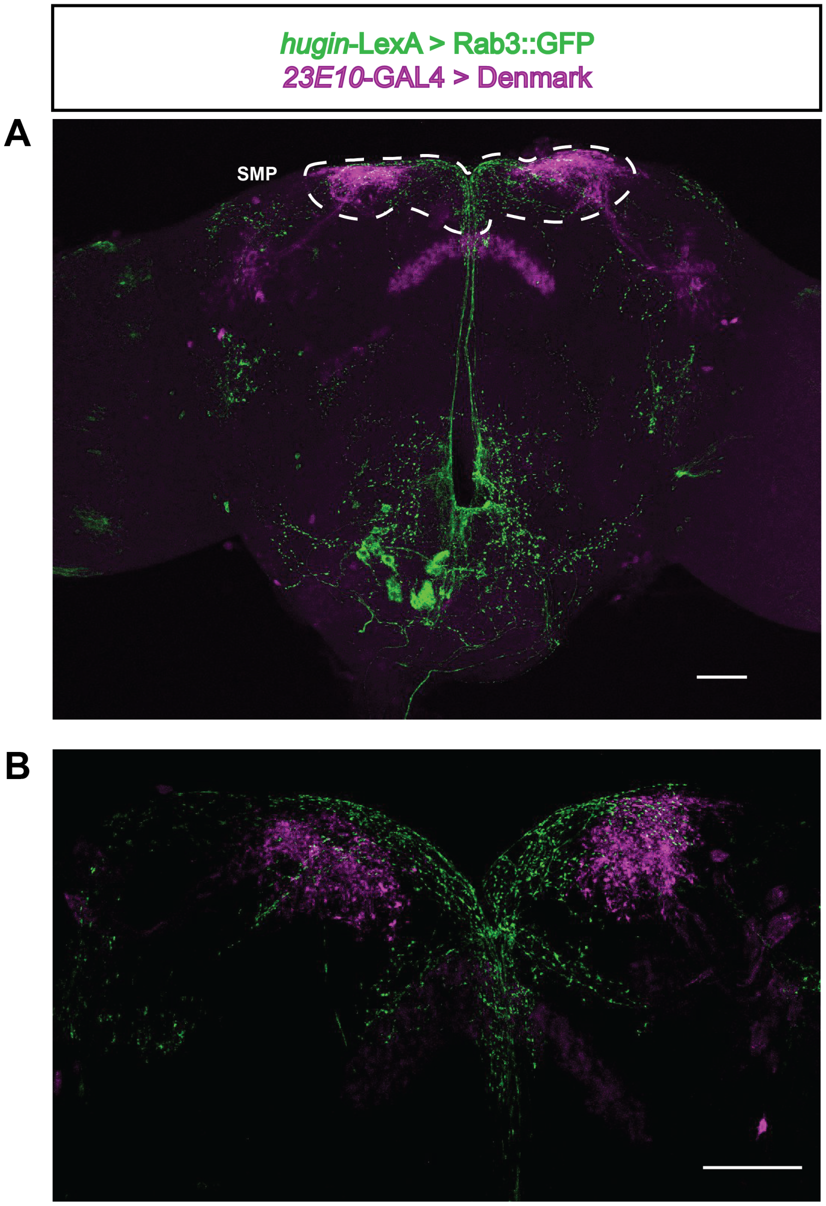


**Figure S2: Presynaptic Sites of *hugin*+ Neurons and *23E10*+ Neuron Dendrites are Located in the SMP.**

Co-labeling of *hugin*+ with RAB3::GFP, a presynaptic marker (green), and 23E10+ neurons with Denmark, a dendritic marker (magenta), and. **(A)** Co-labeling of *hugin*+ and *23E10*+ neurons in the whole fly brain; Superior medial protocerebrum (SMP) region is labeled. **(B)** The SMP region where *hugin*+ projections intermingle with *23E10*+ projections. Scale bars, A: 50 μm; B: 25 μm.

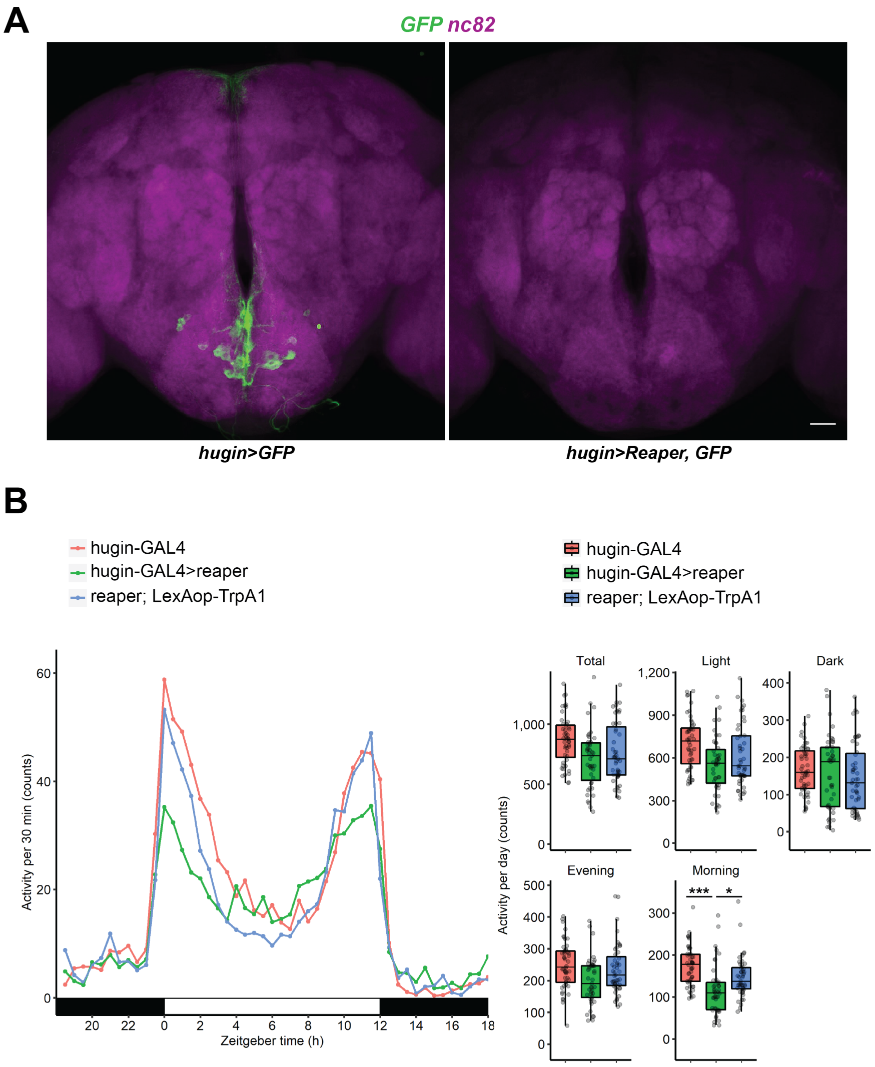


**Figure S3: Expressing Reaper in *hugin*+ Neurons Eliminates *hugin*+ Neurons and Affects the Daytime Activity Profile**

**(A)** Co-labeling of nc82 (magenta) and GFP expressed in *hugin*+ neurons (green). *hugin*+ neurons are absent when Reaper is expressed (right). Scale bar: 25 μm **(B)** Expressing *reaper* in *hugin*+ neurons decreases the morning activity peak. N=43-48. Means compared with one-way ANOVA and Tukey's test. *P < 0.05, **P < 0.01, ***P < 0.001 by Tukey's test after one-way ANOVA.

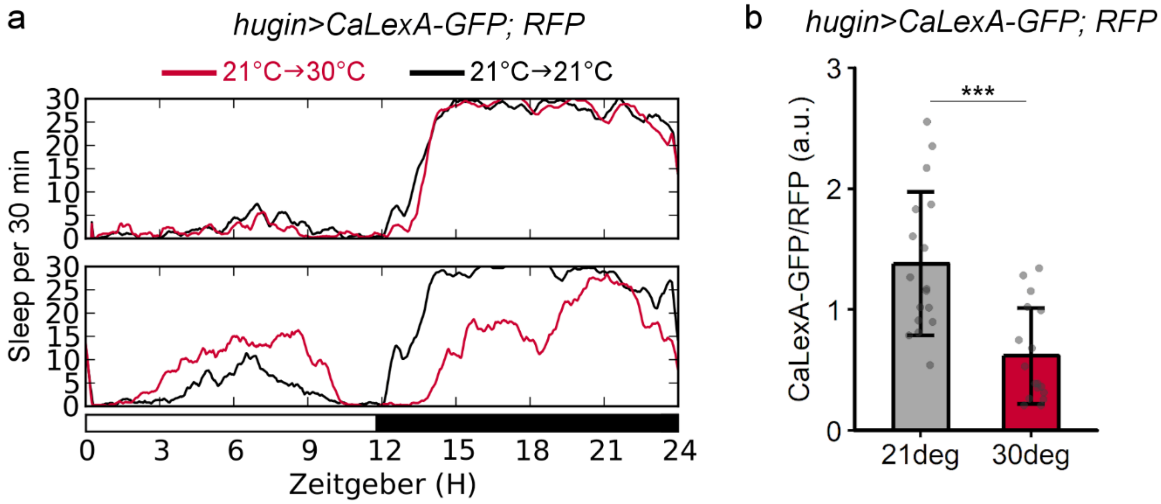


**Figure S4: $Ca^{2+}$ levels of *hugin*+ neurons are suppressed after heat-induced sleep loss.** **(A)** Sleep profiles of *hugin>CaLexA-GFP; RFP* flies at a constant 21°C (black) or at 21°C on the first day (top) and then at 30°C on the second day (bottom, red). The higher temperature on the second day increases daytime sleep and decreases nighttime sleep relative to baseline. **(B)** Tukey's boxplot comparing relative levels of GFP signal normalized to RFP signal in *hugin*+ cell bodies from 21°C control (*n* = 17 flies) or 30°C group (*n* = 17 flies). ***P = 0.000145, Welch's *t*-test.
